# The FHA domain is essential for the autoinhibition of KIF1A/UNC-104

**DOI:** 10.1101/2023.12.24.573241

**Authors:** Shinsuke Niwa, Taisei Watanabe, Kyoko Chiba

**Affiliations:** Frontier Research Institute for Interdisciplinary Sciences (FRIS), Tohoku University, Aramaki-Aoba 6-3, Aoba-ku, Sendai, Miyagi 980-8578, Japan; Department of Biology, Faculty of Science, Tohoku University, Aramaki-Aoba 6-3, Aoba-ku, Sendai, Miyagi 980-8578, Japan

**Keywords:** UNC-104, KIF1A, kinesin-3, axonal transport

## Abstract

KIF1A/UNC-104, a member of the kinesin super family motor proteins, plays a pivotal role in the axonal transport of synaptic vesicles and their precursors. *Drosophila melanogaster* UNC-104 (DmUNC-104) is a relatively recently discovered *Drosophila* kinesin. Although some point mutations that disrupt synapse formation have been identified, the biochemical properties of DmUNC-104 protein have not been investigated. Here, we prepared recombinant full-length DmUNC-104 protein and determined the biochemical features. We analyzed the effect of a previously identified missense mutation in the FHA domain, called *bristly* (*bris*), and found that wild-type DmUNC-104 is a mixture of monomer and dimer and the *bris* mutation strongly promote the dimerization. Furthermore, we found that disease-associated mutations also promote the dimerization of DmUNC-104. These findings suggest that the FHA domain is essential for the autoinhibition of KIF1A/UNC-104, and that abnormal dimerization of KIF1A is linked to human diseases.

## Introduction

Various cellular processes such as cell division, morphogenesis and synapse formation depend on molecular motors. Kinesins, also known as kinesin superfamily proteins (KIFs), are microtubule (MT) -dependent molecular motors that play crucial roles in intracellular transport and cell division (1). Kinesins are classified into 14 families based on their structural and functional features (2). The kinesin-3 family includes eight KIFs in mammals. KIF1A, a member of the kinesin-3 family, is a plus-end directed kinesin specifically expressed in neurons (3). KIF1A and UNC-104, its homolog in *Caenorhabditis elegans* (*C. elegans*) and *Drosophila melanogaster*, are involved in the axonal transport of various cargoes, such as presynaptic components, dense core vesicles (DCVs) and lysosomes (4–8). Genetic variants of *KIF1A* gene cause congenital genetic disorders known as KIF1A-associated neurological disorder (KAND) (9, 10). We have shown that both hyperactivation and impaired activity of KIF1A are associated with neurodegeneration (11–13). Proper control of KIF1A activity is crucial for maintaining neuronal function, however, the molecular mechanisms that regulate KIF1A remain poorly understood.

Most kinesins are dimeric, with the two motor domains (heads) are connected by a coiled-coil domain. Dimeric kinesins alternately use the two heads for processive movement on MTs (14). The kinesin-3 family members have been supposed to be unusual monomeric kinesins as the founding member of kinesin-3, KIF1Bβ and KIF1A, were both described as monomers (3, 15). Later studies showed that *C. elegans* UNC-104 (CeUNC-104) and other kinesin-3 member can form dimers on the cargo membranes (16–18). Moreover, KIF1A has been shown to form dimers in the cytosol (19). Several studies have proposed that kinesin-3 undergoes a monomer to dimer conversion to be activated (20–24). Intramolecular interaction of kinesin-3 mediates its autoinhibition, maintaining the motor in a monomeric state (20, 23, 24). Upon release from autoinhibition, kinesin-3 undergoes dimerization, resulting in increased binding and processivity on MTs compared to the monomeric state (23, 25). Beyond affecting motility, kinesin-3 dimerization plays a significant role in regulating cargo transport, such as DCVs (26). However, the crucial process to initiate the dimerization remains elusive.

As the potential regulator of kinesin-3 dimerization, the autoinhibition of kinesin-3 has been extensively studied. Kinesin-3 members typically possess an N-terminal motor domain (MD) followed by a neck coiled-coil (NC), coiled-coil domain 1 (CC1), FHA domain (FHA), coiled-coil domain 2 (CC2). An early cryo-EM study using CeUNC-104 proposed that CC1 folds back and associates with NC to prevent dimerization (20). Consistent results were shown in subsequent X-ray crystallography using truncated version of KIF1A or KIF13B (22, 23). They showed that MD-NC or CC1-FHA can form dimers, whereas MD-NC-CC1 predominantly forms monomer (22, 23). CC1 was demonstrated to interact with NC and MD, negatively regulating the motor activity of kinesin-3 (23). In addition to MD, NC and CC1, intramolecular interaction between FHA and CC2 have been implicated in the autoinhibition and motor multimerization of KIF1A (21). Those analyses have mainly used only the respective domains, and the role of these domains on the regulation of kinesin-3, especially in the context of full-length protein, are not well understood.

*Drosophila melanogaster* has been an important model organism in the study of the kinesin superfamily (27, 28). *Drosophila* kinesin-1, a homolog of KIF5, has been well-known since the early stages of kinesin research and has been extensively analyzed. The fundamental biochemical properties of kinesin-1 were determined using the *Drosophila* kinesin-1 protein (29–32). Compared to *Drosophila* kinesin-1, *Drosophila melanogaster* UNC-104 (DmUNC-104) (also known as *Drosophila* KIF1A or IMAC) is a relatively recently discovered member of *Drosophila* kinesins (33). In addition to nonsense mutations, some point mutations that disrupt synapse formation has been identified (34, 35). Even though measuring the biochemical changes caused by those point mutations may help us understand the basic properties of kinesin-3, the biochemical properties of the DmUNC-104 protein have not been investigated.

Here, we purified and analyzed the activity of full-length DmUNC-104 protein. We show that mutations previously identified in or near the FHA domain induce the dimerization of DmUNC-104. These mutations nearly abolished monomer formation, suggesting they initiated the complete release of DmUNC-104 autoinhibition. In single molecule assays using a total internal reflection fluorescent (TIRF) microscope, the dimerized mutant DmUNC-104 showed elevated activity compared to the monomeric wild type. Collectively, our data suggest that the FHA domain plays an essential role in the autoinhibition of KIF1A/UNC-104.

## Results

### The full-length DmUNC-104 exists in monomeric and dimeric forms

Full-length DmUNC-104, tagged with a C-terminal superfolder GFP (sfGFP) and Strep-tag II (DmUNC-104^WT^, calculated M.W. = 220 kDa), was expressed using the baculovirus expression system (Fig. 1A). Following affinity purification, we examined the oligomeric state of the purified protein through SEC and mass photometry. DmUNC-104^WT^ displayed two distinct peaks upon elution (Fig. 1B). An additional peak near the void volume was observed, but no considerable amount of DmUNC-104^WT^ was detected in this peak. Upon subjecting the peak to agarose gel electrophoresis, bands corresponding to the size of insect ribosomal RNA (∼2 and 4 kbp) were observed (data not shown). The bands disappeared by RNaseI treatment (data not shown), confirming that the additional peak mainly contains contaminated RNA. Therefore, we focused our analysis on the latter two peaks. Mass photometry analysis at 10 nM (concentration assumed as dimers) confirmed that both peaks contained monomers and dimers. The earlier peak (peak-1) consists of particles corresponding to 209 ± 42 kDa and 414 ± 39 kDa, accounting for 24% and 54% of the population, respectively (Fig. 1C). The later peak (peak-2) contained particles corresponding to 216 ± 18 kDa and 435 ± 31 kDa with 71% and 10% of the populations, respectively (Fig. 1C). The observed molecular weights align with a monomer and a dimer of DmUNC-104^WT^, considering the calculated mass of 220 kDa as a monomer. These results indicate that full-length DmUNC-104 can exist as both monomers and dimers, and the state might be in equilibrium, as both monomers and dimers are present in individual peaks even separation by SEC.

**Figure 1.**
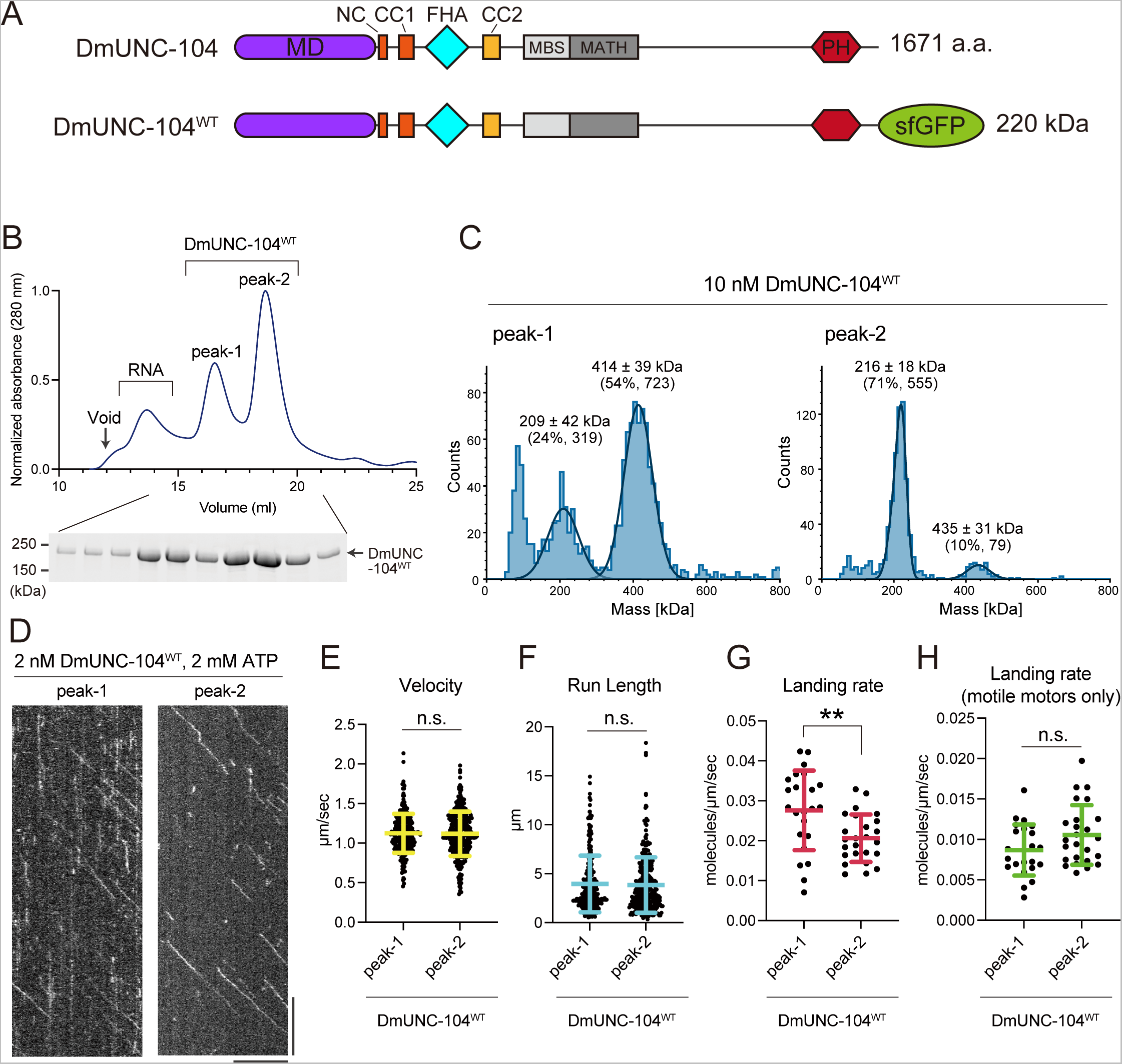
The full-length DmUNC-104 exists in monomeric and dimeric forms. **(A)** Schematic diagram of the domain architecture of *Drosophila* UNC-104 (DmUNC-104) and the DmUNC-104 fused with sfGFP and 2xStrep-tag II (DmUNC-104^WT^). MD: motor domain, NC: Neck coiled-coil, CC1: Coiled-coil 1 domain, FHA: forkhead-associated domain, CC2: Coiled-coil 2 domain, MBS: membrane-associated guanylate kinase homolog (MAGUK)-binding stalk domain, MATH: meprin and TRAF homology domain, PH: Pleckstrin homology domain, sfGFP: superfolder green fluorescent protein. Calculated molecular weight (M.W.) of each is shown. **(B)** Size exclusion chromatography of DmUNC-104^WT^. The void volume of the column and the RNA-rich peak are indicated. SDS-PAGE of the elution fractions is shown beneath the chromatogram. Numbers shown at the left side of the gel indicate molecular weight standard. **(C)** Mass photometry of DmUNC-104^WT, peak-1^ and DmUNC-104^WT, peak-2^. Histogram shows the particle count at 10 nM along the indicated molecular weight. The concentration assumes the protein in its dimeric form. The lines show Gaussian fits (mean ± standard deviation (S.D.). Percentage of the particle in the peak and particle counts are indicated. **(D)** Representative kymograph showing the motility of DmUNC-104^WT, peak-1^ and DmUNC-104^WT, peak-2^ at 2 nM in the presence of 2 mM ATP. The concentration assumes the protein in its dimeric form. Horizontal and vertical bars show 10 μm and 10 seconds, respectively. **(E)** Dot plots showing the velocity of DmUNC-104^WT, peak-1^ and DmUNC-104^WT, peak-2^. Each dot shows a velocity of a molecule. Green bars represent mean ± S.D.. n = 260 and 312 for peak-1 and peak-2, respectively. ns, no statistically significant difference in t-test. **(F)** Dot plots showing the run length of DmUNC-104^WT, peak-1^ and DmUNC-104^WT, peak-2^. The distance each molecule traveled along microtubule was measured. Each dot shows a run length of a molecule. Green bars represent median value and interquartile range. n = 260 and 312 for peak-1 and peak-2, respectively. ns, no statistically significant difference in Mann-Whitney U test. **(G)** Dot plots showing the landing rate of DmUNC-104^WT, peak-1^ and DmUNC-104^WT, peak-2^. The number of molecules binding per 1-µm of microtubules per second is shown. Each dot shows a single datum point. Green bars represent median value and interquartile range. n = 23 and 25 for peak-1 and peak-2, respectively. Mann-Whitney U test. **, p < 0.01 (p value = 0.0040). **(H)** Dot plots showing the landing rate of motile motors in DmUNC-104^WT, peak-1^ and DmUNC-104^WT, peak-2^. Molecules showing processive motion on microtubules only were counted. The number of motile motors per 1-µm of microtubules per second is shown. Each dot shows a single datum point. Green bars represent median value and interquartile range. n = 23 and 25 for peak-1 and peak-2, respectively. ns, no statistically significant difference in Mann-Whitney U test.

### The motilities of the two peaks were overall indistinguishable

We compared the motility of peak-1 and peak-2 of DmUNC-104^WT^ by single molecule assay using a TIRF microscope. Both peaks were examined at 2 nM, at the concentration assumed as dimers (equivalent to 4 nM of monomer). Peak-1 and peak-2 showed almost identical properties on the single molecule assays (Fig. 1D). The velocity was comparable (1.12 ± 0.25 µm/s and 1.12 ± 0.28 µm/s, mean ± S.D. for peak-1 and peak-2, respectively) and the run length showed no significant difference (2.99 µm and 3.07 µm, median values for peak-1 and peak-2, respectively) (Fig. 1E and 1F). When we calculated landing rates of the motors on the microtubules, peak-1 showed a higher landing rate than peak-2 (0.028 and 0.020 molecules/µm/s, median values for peak-1 and peak-2, respectively) (Fig. 1G). However, no significance was observed in the landing rates when considering motile molecules out of the total landings, which include both motile and immotile landings (0.0087 and 0.0099 molecules/µm/s, median values for peak-1 and peak-2, respectively) (Fig. 1H). These results suggest that peak-1 and peak-2 contain a similar number of motile motors, but peak-1 contains more immotile motors that can bind to microtubules without processive motion.

### The *bris* mutation induces DmUNC-104 dimerization and activation

The *bristly* (*bris*) mutation was isolated through EMS mutagenesis in *Drosophila*, leading to an increased number of dendritic filopodia (36). Subsequent studies identified *bris* as an allele of *unc-104*, with the responsible mutation being R561H (R562H in the isoform we utilized, hereafter referred to as R562H) (35). The study also revealed that *bris* causes defective synaptogenesis, possibly due to impaired axonal transport of presynaptic materials such as Bruchpilot (Brp). The R562 residue is located in the β11-loop of the FHA domain of DmUNC-104. To investigate the effect of the *bris* mutation on the biochemical properties of DmUNC-104, we introduced the corresponding R562H mutation into DmUNC-104. Firstly, we examined the dimerization state (Fig. 2A). Unlike DmUNC-104^wt^, DmUNC-104*^bris^* was eluted as a single peak in SEC (Fig. 2B), with the elution volume similar to that of peak-1 of DmUNC-104^wt^ protein. Mass photometry of the peak revealed particles corresponding to 237 ± 85 kDa and 476 ± 58 kDa, accounting for 12% and 71% of the population, respectively (Fig. 2C). This confirmed that DmUNC-104*^bris^* contains a small number of monomers but predominantly forms dimers. In single molecule assays, DmUNC-104*^bris^*showed a higher velocity than that of peak-1 of wild-type (1.12 ± 0.25 µm/s and 1.25 ± 0.27 µm/s, mean ± S.D. for DmUNC-104^WT, peak-1^ and DmUNC-104*^bris^*, respectively) (Fig. 2D and 2E). No significant differences were observed in the run length (2.99 µm and 3.45 µm, median values for DmUNC-104^WT, peak-1^ and DmUNC-104*^bris^*, respectively) compared with peak-1 of wild-type (Fig. 2F). However, we observed higher landing rates in both total landing (0.028 and 0.034 molecules/µm/s, median values for DmUNC-104^WT, peak-1^ and DmUNC-104*^bris^*, respectively) and motile motors’ landing (0.0087 and 0.015 molecules/µm/s, median values for DmUNC-104^WT, peak-1^ and DmUNC-104*^bris^*, respectively) (Fig. 2G and 2H). These suggest that DmUNC-104*^bris^* is biochemically more active than DmUNC-104^WT^.

**Figure 2.**
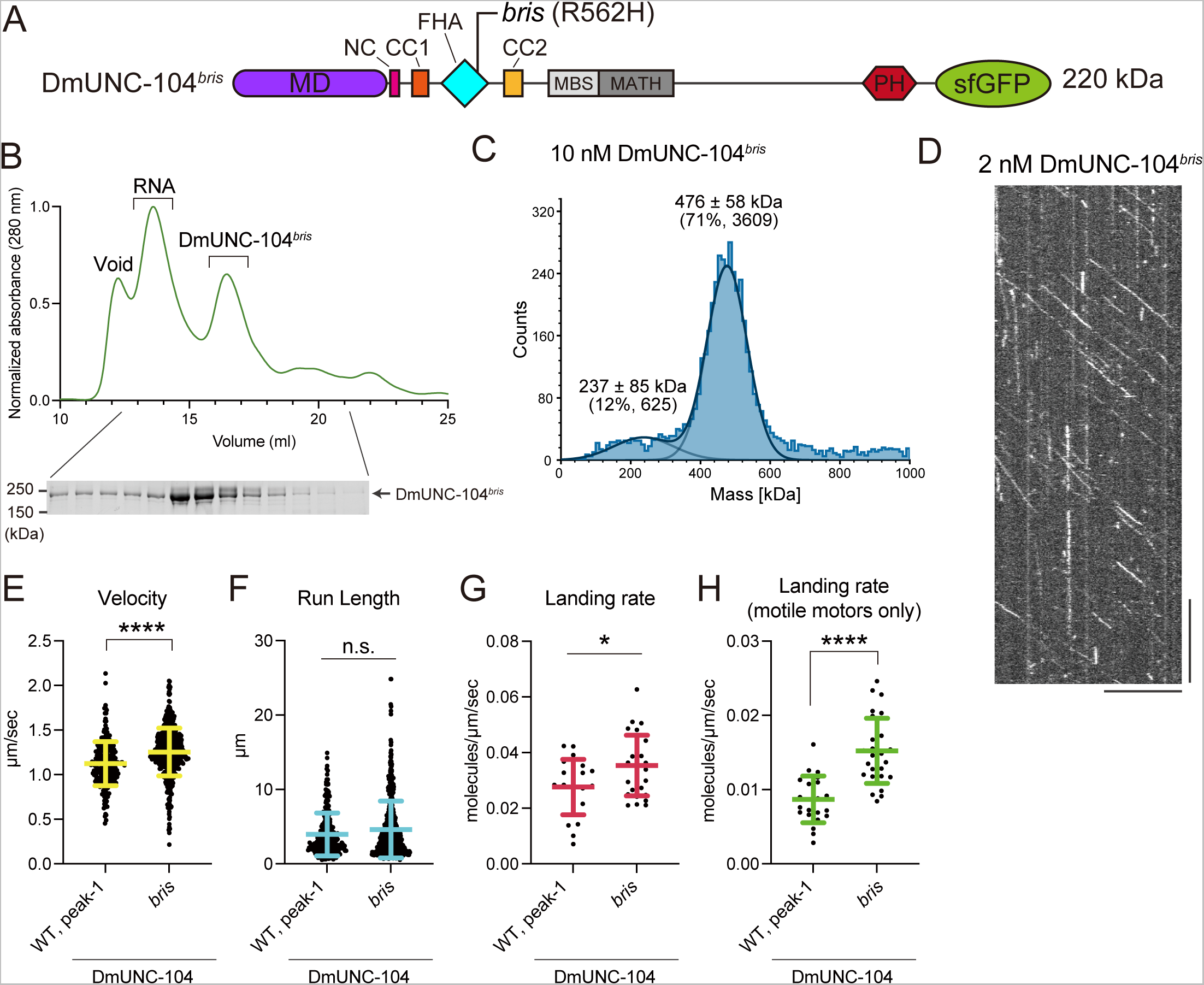
The *bris* mutation induces DmUNC-104 dimerization. **(A)** Schematic drawing of the domain organization of DmUNC-104*^bris^*. Calculated molecular weight (M.W.) is shown. **(B)** Size exclusion chromatography of DmUNC-104*^bris^*. The void volume of the column and the RNA-rich peak are indicated. SDS-PAGE of the elution fractions is shown beneath the chromatogram. Numbers shown at the left side of the gel indicate molecular weight standard. **(C)** Mass photometry of DmUNC-104*^bris^*. Histogram shows the particle count at 10 nM along the indicated molecular weight. The concentration assumes the protein in its dimeric form. The lines show Gaussian fits (mean ± standard deviation (S.D.). Percentage of the particle in the peak and particle counts are indicated. **(D)** Representative kymograph showing the motility of 2 nM DmUNC-104*^bris^* in the presence of 2 mM ATP. The concentration assumes the protein in its dimeric form. Horizontal and vertical bars show 10 μm and 10 seconds, respectively. **(E)** Dot plots showing the velocity of DmUNC-104^WT, peak-1^ and DmUNC-104*^bris^*. Each dot shows a velocity of a molecule. Green bars represent mean ± S.D.. n = 260 and 582 for DmUNC-104^WT, peak-1^ and DmUNC-104*^bris^*, respectively. t-test. ****, p < 0.0001. DmUNC-104^WT, peak-1^ values are replotted from Fig. 1E. **(F)** Dot plots showing the run length of DmUNC-104^WT, peak-1^ and DmUNC-104*^bris^*. The distance each molecule traveled along microtubule was measured. Each dot shows a run length of a molecule. Green bars represent median value and interquartile range. n = 260 and 582 for DmUNC-104^WT, peak-1^ and DmUNC-104*^bris^*, respectively. ns, no statistically significant difference in Mann-Whitney U test. DmUNC-104^WT, peak-1^ values are replotted from Fig. 1F. **(G)** Dot plots showing the landing rate of DmUNC-104^WT, peak-1^ and DmUNC-104*^bris^*. The number of molecules binding per 1-µm of microtubules per second is shown. Each dot shows a single datum point. Green bars represent median value and interquartile range. n = 23 and 28 for DmUNC-104^WT, peak-1^ and DmUNC-104*^bris^*, respectively. Mann-Whitney U test. *, p < 0.05 (p value = 0.0392). DmUNC-104^WT, peak-1^ values are replotted from Fig. 1G. **(H)** Dot plots showing the motile motors’ landing rate of DmUNC-104^WT, peak-1^ and DmUNC-104*^bris^*. Molecules showing processive motion on microtubules only were counted. The number of motile motors per 1-µm of microtubules per second is shown. Each dot shows a single datum point. Green bars represent median value and interquartile range. n = 23 and 28 for DmUNC-104^WT, peak-1^ and DmUNC-104*^bris^*, respectively. Mann-Whitney U test. ****, p < 0.0001. DmUNC-104^WT, peak-1^ values are replotted from Fig. 1H.

### The *bris* mutation causes gain of *unc-104* function in *C. elegans* neuron

The previous study suggested that the *bris* mutation is a putative loss-of-function mutation, as the phenotype was exacerbated in transheterozygotes with a null allele (35). Our results from SEC and single molecule assays showed that the *bris* mutation causes dimerization and activation of DmUNC-104. These results imply that the *bris* mutation might be a gain-of-function mutation. We have shown that *C. elegans* is a good model to test whether a kinesin-3 mutation is loss-of-function or gain-of-function in the axonal transport of synaptic materials. This assay takes advantage of the fact that axonal transport of synaptic vesicles is reduced in *arl-8* mutants (37). If the *unc-104* mutation is gain-of-function, it works as a suppressor of the *arl-8* phenotype, whereas if the *unc-104* mutation is loss-of-function, it works as an enhancer of the *arl-8* phenotype. Using this system, we analyzed the *bris* mutation. R562 in DmUNC-104 corresponds to R551 in CeUNC-104 (Fig. 3A). We introduced the *bris* mutation in *C. elegans unc-104* gene by CRISPR/cas9 (38, 39) (Supplementary Figure S1). We observed the phenotype of the DA9 neuron. DA9 neuron is a polarized neuron with distinct regions, including a dendrite, cell body and axon (40–44) (Fig. 3B). En passant synapses are formed along the axon of the DA9 neuron, visualized by a synaptic vesicle (SV) marker, GFP::RAB-3 (Fig. 3B and 3C). In *arl-8* worms lacking ARL-8, due to the reduced axonal transport, SVs mislocalized to the commissure and proximal asynaptic region (Fig. 3D). In *arl-8; unc-104^R551H^* double mutant worms, however, the localization of RAB-3 signals was partially restored (Fig. 3E). To confirm the phenotype, we counted the number of GFP::RAB-3 puncta in the commissure (Fig. 3F) and measured the length of asynaptic region (Fig. 3G). The aberrant localization of synaptic vesicles in *arl-8* worms was suppressed by *unc-104*^R551H^ mutation. Moreover, we found *unc-104^R551H^* could suppress the phenotype of *arl-8* in an autosomal dominant manner. These results indicate that the *bris* mutation in *unc-104* might be a gain-of-function mutation, not a loss-of-function mutation.

**Figure 3.**
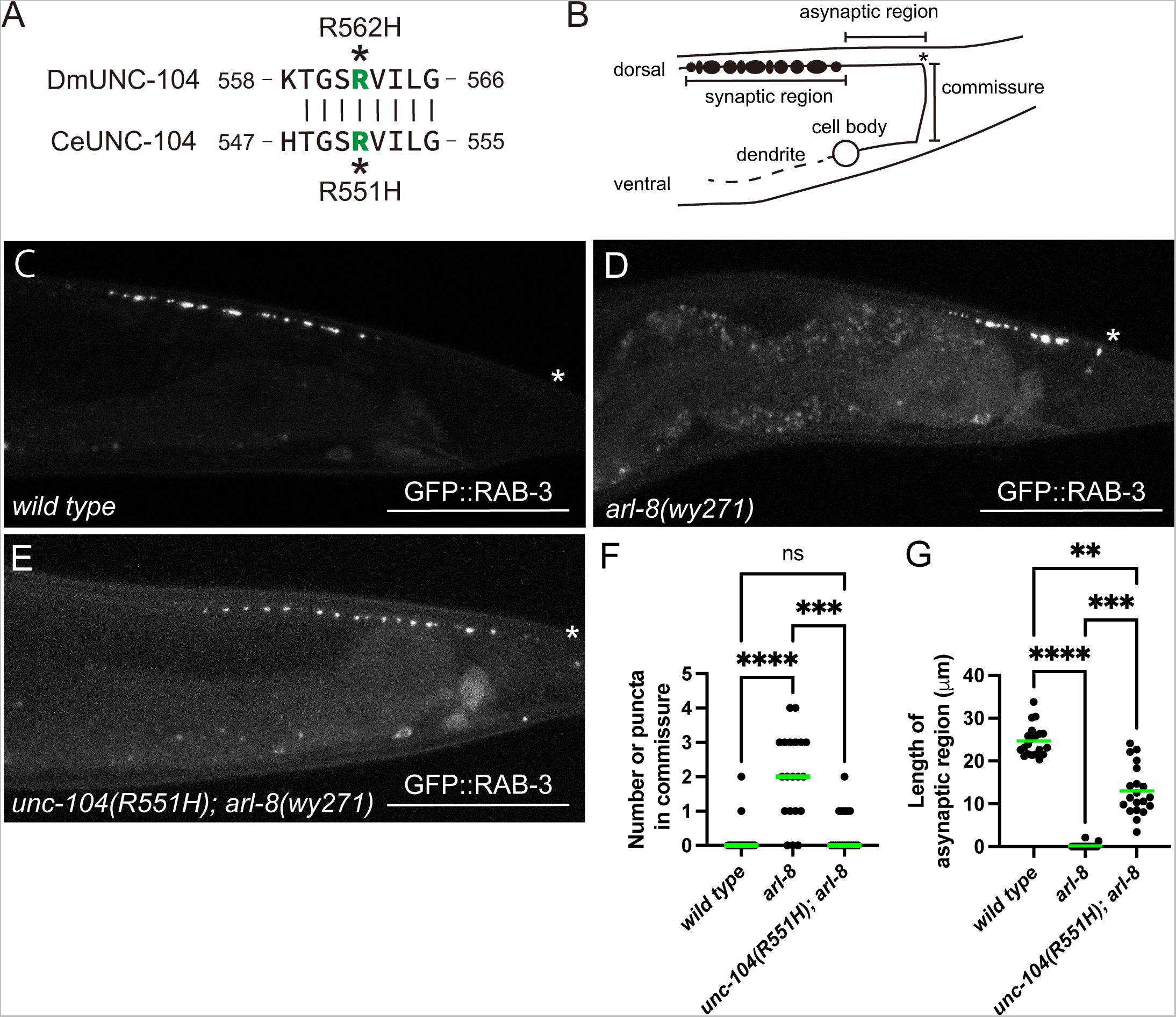
The *bris* mutation causes gain of *unc-104* function in *C. elegans* neuron. **(A)** Sequence comparison between *Drosophila* UNC-104 and *C. elegans* UNC-104. R562H in *Drosophila* UNC-104 corresponds to R551 in *C. elegans* UNC-104. **(B)** Schematic drawing of the DA9 neuron of *C. elegans*, showing the cell body, dendrite, commissure, dorsal synaptic region and asynaptic region. The asterisk indicates the location where the commissure joins the dorsal nerve cord. Puncta show the localization of the synaptic vesicles or their precursors represented by GFP::RAB-3. **(C-E)** GFP::RAB-3 was expressed in DA9 neuron under the *itr-1* promoter. Representative images showing the localization of GFP::RAB-3 in respective worms. **(C)** wild type. **(D)** *arl-8*(wy271). **(E)** *arl-8*(wy271);*unc-104*(R551H). Integrated marker *wyIs85* [*Pitr-1::GFP::RAB-3*] was used. Asterisks indicate the commissure bend shown in A. Bars, 50 μm. **(F)** The length of asynaptic region. n = 20. 1-way ANOVA followed by Tukey’s multiple comparisons test. ns, no statistically significant difference; ***, p < 0.001; ****, p < 0.0001. **(G)** The number of RAB-3 puncta misaccumulated in the commissure. n = 20. Kruskal–Wallis 1-way ANOVA on ranks and Dunn’s multiple comparisons test. **, p < 0.01; ***, p < 0.001; ****, p < 0.0001.

### DmUNC-104 is converted to a dimer upon autoinhibition release

A previous study has shown that the FHA and CC2 domains form intramolecular interactions by bending at the hinge sequence “QGID”, which is well conserved across species (Fig. 4A). It has been suggested that a mutation in the hinge disrupts the association between the FHA and CC2 domains of human KIF1A (21). Consistent with this, a mutation in the hinge works as a suppressor of the *arl-8* phenotype in *C. elegans* (42). To confirm that the effect of the *bris* mutation is caused by the release from the autoinhibition, we introduced the hinge mutation, G618R mutation, into DmUNC-104 and analyzed its biochemical behavior (Fig. 4A and 4B). Similar to DmUNC-104*^bris^*, DmUNC-104^G618R^ was eluted as a single peak in SEC, with the elution volume corresponding to peak-1 of DmUNC-104^WT^ (Fig. 4C). Mass photometry of the peak revealed particles distributed almost in a single peak, with a molecular weight of 437 ± 70 kDa, accounting for 80% of the populations (Fig. 4D). This molecular weight corresponds to the dimer of DmUNC-104^G618R^. We further tested the motility of DmUNC-104^G618R^ by single molecule assay (Fig. 4E). Compared with peak-1 of DmUNC-104^WT^, DmUNC-104^G618R^ showed slightly higher velocity (1.12 ± 0.25 µm/s and 1.21 ± 0.24 µm/s, mean ± S.D. for DmUNC-104^WT, peak-1^ and DmUNC-104^G618R^, respectively) (Fig. 4F). No significant differences were observed in the run length (2.99 µm and 3.15 µm, median values for DmUNC-104^WT, peak-1^ and DmUNC-104^G618R^, respectively) and the landing rate (0.028 and 0.030 molecules/µm/s, median values for DmUNC-104^WT, peak-1^ and DmUNC-104^G618R^, respectively) compared with peak-1 of wild-type (Fig. 4G and 4H). However, we observed higher landing rates in motile motors’ landing (0.0087 and 0.012 molecules/µm/s, median values for DmUNC-104^WT, peak-1^ and DmUNC-104^G618R^, respectively) (Fig. 4I). These properties observed in DmUNC-104^G618R^ are similar to those of DmUNC-104*^bris^*, suggesting that the *bris* mutation disrupt the autoinhibition of DmUNC-104.

**Figure 4.**
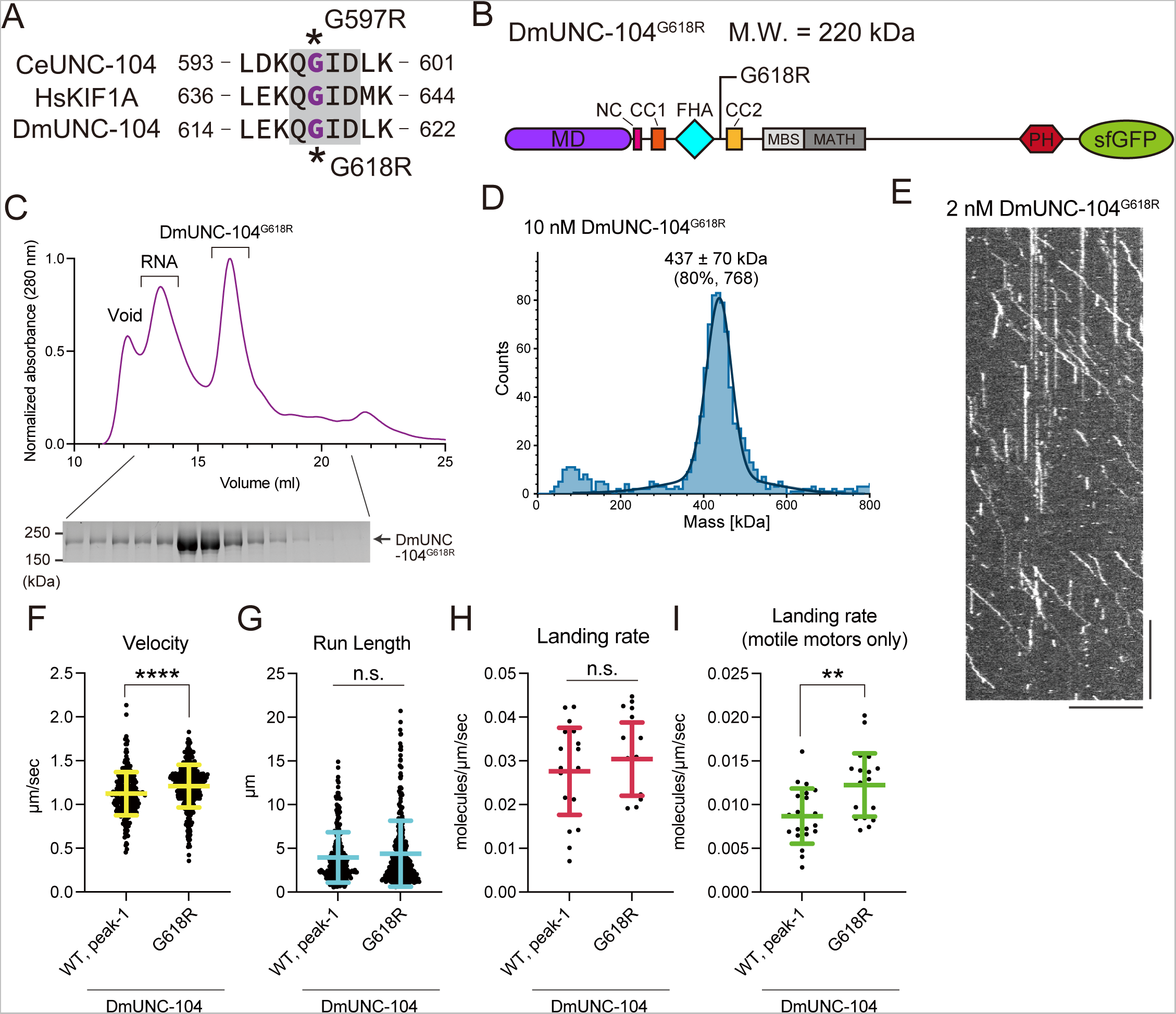
DmUNC-104 is converted to a dimer upon autoinhibition release. **(A)** Sequence alignment of *C. elegans* UNC-104, human KIF1A and *Drosophila* UNC-104. The hinge sequence “QGID” between FHA and CC2 was highlighted in gray. G597 in *C. elegans* UNC-104 corresponds to G618 in *Drosophila* UNC-104. **(B)** Schematic drawing of the domain organization of DmUNC-104^G618R^. Calculated molecular weight (M.W.) is shown. **(C)** Size exclusion chromatography of DmUNC-104^G618R^. The void volume of the column and the RNA-rich peak are indicated. SDS-PAGE of the elution fractions is shown beneath the chromatogram. Numbers shown at the left side of the gel indicate molecular weight standard. **(D)** Mass photometry of DmUNC-104^G618R^. Histogram shows the particle count 10 nM along the indicated molecular weight. The concentration assumes the protein in its dimeric form. The lines show Gaussian fits (mean ± standard deviation (S.D.). Percentage of the particle in the peak and particle counts are indicated. **(E)** Representative kymograph showing the motility of 2 nM DmUNC-104^G618R^ in the presence of 2 mM ATP. The concentration assumes the protein in its dimeric form. Horizontal and vertical bars show 10 μm and 10 seconds, respectively. **(F)** Dot plots showing the velocity of DmUNC-104^WT, peak-1^ and DmUNC-104^G618R^. Each dot shows a velocity of a molecule. Green bars represent mean ± S.D.. n = 260 and 354 for DmUNC-104^WT, peak-1^ and DmUNC-104^G618R^, respectively. t-test. ****, p < 0.0001. DmUNC-104^WT, peak-1^ values are replotted from Fig. 1E. **(G)** Dot plots showing the run length of DmUNC-104^WT, peak-1^ and DmUNC-104^G618R^. The distance each molecule traveled along microtubule was measured. Each dot shows a run length of a molecule. Green bars represent median value and interquartile range. n = 260 and 354 for DmUNC-104^WT, peak-1^ and DmUNC-104^G618R^, respectively. ns, no statistically significant difference in Mann-Whitney U test. DmUNC-104^WT, peak-1^ values are replotted from Fig. 1F. **(H)** Dot plots showing the landing rate of DmUNC-104^WT, peak-1^ and DmUNC-104^G618R^. The number of molecules binding per 1-µm of microtubules per second is shown. Each dot shows a single datum point. Green bars represent median value and interquartile range. n = 23 and 22 for DmUNC-104^WT, peak-1^ and DmUNC-104^G618R^, respectively. ns, no statistically significant difference in Mann-Whitney U test. DmUNC-104^WT, peak-1^ values are replotted from Fig. 1G. **(I)** Dot plots showing the motile motors’ landing rate of DmUNC-104^WT, peak-1^ and DmUNC-104^G618R^. Molecules showing processive motion on microtubules only were counted. The number of motile motors per 1-µm of microtubules per second is shown. Each dot shows a single datum point. Green bars represent median value and interquartile range. n = 23 and 22 for DmUNC-104^WT, peak-1^ and DmUNC-104^G618R^, respectively. Mann-Whitney U test. **, p < 0.01 (p value = 0.0011). DmUNC-104^WT, peak-1^ values are replotted from Fig. 1H.

### Disease-associated mutations induce the dimerization of DmUNC-104

The data above suggested that DmUNC-104 is a promising protein to study the monomer-to-dimer conversion. Therefore, we investigated whether disease-associated human mutations affect the dimerization of DmUNC-104. While most disease-associated mutations are known to have defective effects on KIF1A (9, 45–47), our previous work suggested that some mutations are gain-of-function mutations for *KIF1A* (12). We have demonstrated that recombinant proteins of KIF1A^V8M^, KIF1A^A255V^ and KIF1A^R350G^ show increased microtubule binding in single molecule assays (12). However, the precise molecular mechanisms of KIF1A activation induced by these mutations remain totally unknown. To study the effect of these disease-associated mutations on the dimerization of DmUNC-104, we introduced corresponding mutations, V6M, A255V and R348G, into DmUNC-104 (Fig. 5A and 5B). Our SEC analysis revealed that DmUNC-104^V6M^ and DmUNC-104^R348G^ predominantly eluted as a single peak, corresponding to peak-1 of the wild-type protein (Fig. 5C). While they showed a small peak around the elution volume corresponding to peak-2 of the wild-type protein, it was not as pronounced. In contrast, DmUNC-104^A255V^ eluted into two peaks similar to DmUNC-104^WT^ (Fig. 5C). These results indicate that among the disease-associated gain-of-function mutations, at least two mutations may induce dimerization of KIF1A/UNC-104. Taken together, our data suggest that the dimerization of KIF1A/UNC-104 is regulated by intramolecular interactions involving the FHA domain, which plays an essential role. Aberrant dimerization may potentially contribute to the development of human diseases.

**Figure 5.**
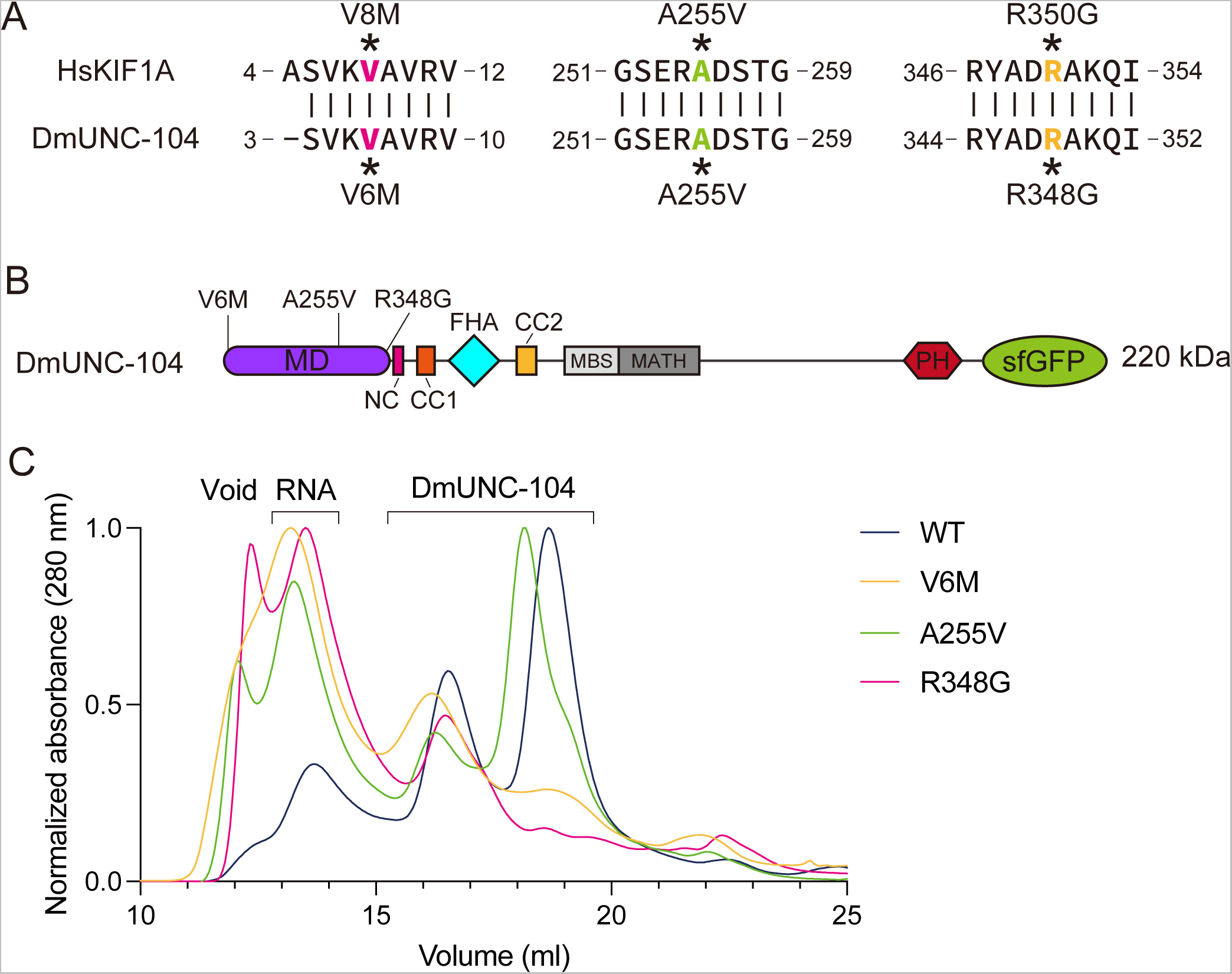
Disease-associated mutations affect dimerization of DmUNC-104. **(A)** Sequence comparison between human KIF1A and *Drosophila* UNC-104. V8, A255 and R350 in human KIF1A correspond to V6, A255 and R348 in *Drosophila* UNC-104. **(B)** Size exclusion chromatography of DmUNC-104^WT^ (blue), DmUNC-104^V6M^ (pink), DmUNC-104^A255V^ (green) and DmUNC-104^R348G^ (orange). DmUNC-104^WT^ values are replotted from Fig. 1B. The void volume of the column and the RNA-rich peak are indicated.

## Discussion

### Autoinhibition release sufficiently induces dimerization in full-length KIF1A/UNC-104

We showed that a deletion mutant of *C. elegans* UNC-104, consisting of the MD-CC1-FHA-CC2 domains, exists in an equilibrium between dimers and monomers (25). However, we could not analyze the property of the full-length *C. elegans* UNC-104 protein due to the difficulty in purification. In this study, we succeeded in purifying full-length DmUNC-104. Using the full-length DmUNC-104 protein, we now clearly showed that autoinhibition release is sufficient to convert monomeric KIF1A/UNC-104 molecule into dimers. Previous studies have reported the interaction between the FHA and CC2 domains (Fig. 6A), suggesting that disruption of this interaction is involved in the activation of KIF1A/UNC-104 (21). However, it remained unclear how this disruption affects the overall conformation of the full-length KIF1A/UNC-104. In this study, we analyzed G618R mutation which disrupts the autoinhibitory interaction between the FHA and CC2 domains. We showed that G618R almost completely converts monomeric DmUNC-104 into dimers. Furthermore, the *bris* mutation, which putatively interferes with the FHA-CC2 interaction, also promoted the dimerization of DmUNC-104. Our data suggest that the interaction between the FHA and CC2 domains is the key regulator for releasing the autoinhibition of KIF1A/UNC-104 (Fig. 6A and 6B).

**Figure 6.**
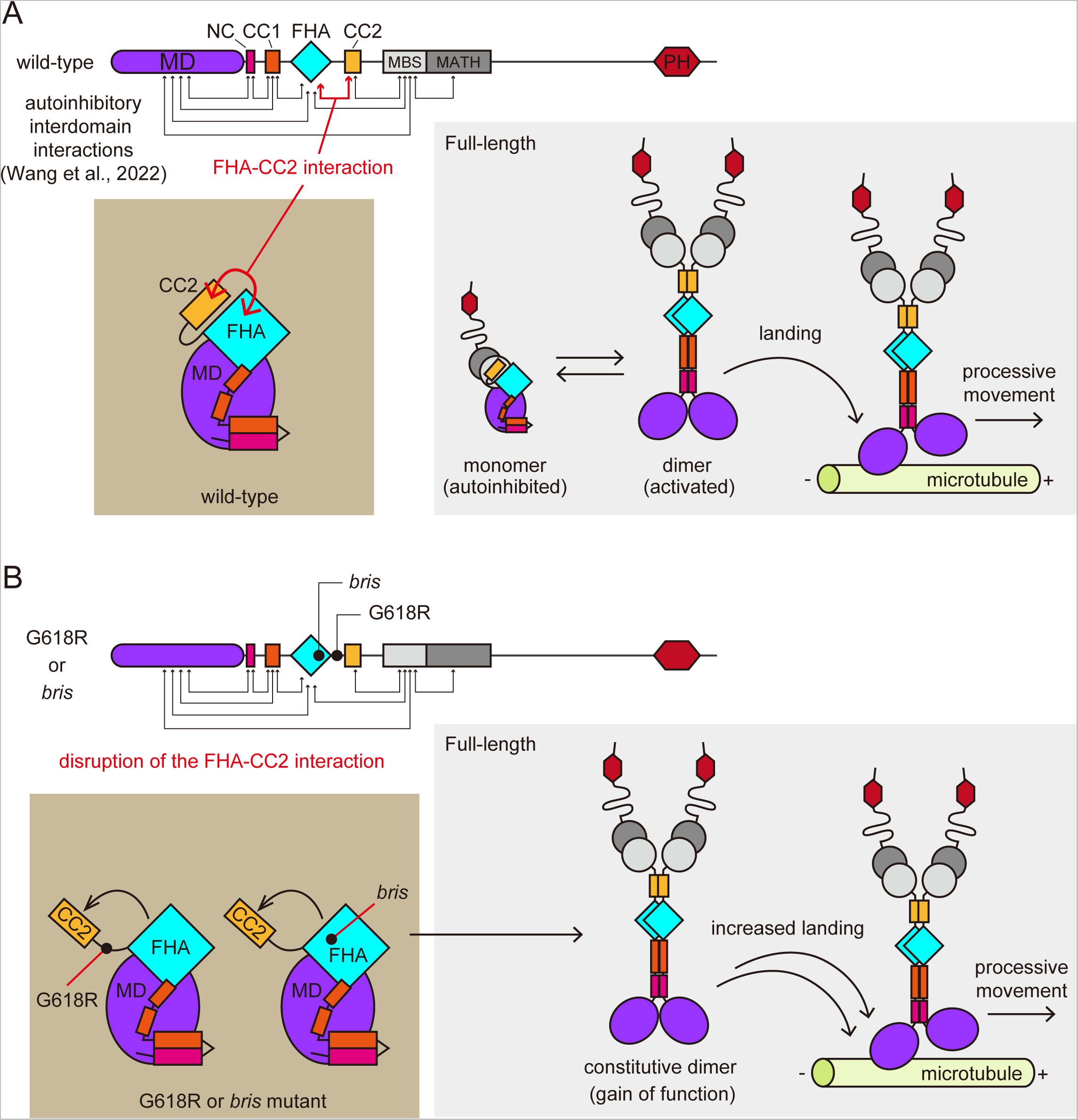
Models for the activation of KIF1A/UNC-104. Schematic models for the autoinhibition release of KIF1A/UNC-104. **(A)** In the wild-type protein, the molecule is autoinhibited by multivalent interdomain interactions, however, due to the equilibrium, dimerization occurs at a certain possibility. The dimerized motor lands on microtubules and start processive movement. **(B)** In G618R or *bris* mutant, the FHA-CC2 interaction is disrupted. The disruption initiates complete release of KIF1A/UNC-104 from the autoinhibition. As a result, G618R or *bris* mutant forms stable dimers. The mutants show higher microtubule-landing rate *in vitro* and gain-of-function phenotype *in vivo*.

### The *bris* mutation disrupts the intramolecular interaction between the FHA and CC2 domains

A recent study revealed the structure of a kinesin-3 member protein, KLP-6 (kinesin-like protein 6 in *C. elegans*), showing that the entire molecule cooperatively self-folds to form an inactive monomer (Fig. 6A) (24). The study showed that the FHA domain interacts not only with CC2 but also with MD and CC1 (Fig. 6A). In the structure, R538 of KLP-6, equivalent to R562 of DmUNC-104, is on the contact site of the FHA domain and CC2 (Supplementary Figure S2). Consistent with the structure, the *bris* mutation induces the dimerization of DmUNC-104. The properties of DmUNC-104*^bris^*are similar to those of the hinge mutant DmUNC-104^G618R^ whose FHA-CC2 interaction is predicted to be disrupted (Fig. 6B).

The *bris* mutation was initially characterized as a loss-of-function mutation through genetic assays (35). However, our biochemical analyses and genetic experiments in *C. elegans* consistently suggest that the *bris* mutation represents a gain-of-function mutation. One possible explanation is that the initially identified *bris* mutant phenotypes might arise from a gain-of-function behavior of the DmUNC-104 molecule. Another possibility is that the effect of the release of the autoinhibition of KIF1A/UNC-104 may vary among species and cell types. It is possible that the *bris* mutation affects the binding of other molecules, including cargos, and the impact of altered interactions may be more severe in *Drosophila*, causing the observed loss-of-function phenotype. This parallels the known case of kinesin-1, where disrupted autoinhibition leads to a loss-of-function phenotype in *Drosophila* (48). To investigate these possibilities, further genetic studies using *Drosophila* will be required.

### KIF1A dimerization and human diseases

In our previous work, we demonstrated that some disease-associated KIF1A mutations, specifically V8M, A255V and R350G, increase the activity of KIF1A (12). However, the precise molecular mechanism has remained to be clarified. In the current study, we found that V8M and R350G mutations induce the dimerization of DmUNC-104. Considering that disrupted autoinhibition, induced by the *bris* and hinge mutations, promoted dimerization of DmUNC-104 (Fig. 2 and 4), these disease-association mutations would release the autoinhibition of KIF1A and enhance the activity. A similar observation was found in an ALS-related mutant of KIF5A, a member of conventional kinesin. The ALS-associated mutations result in aberrant oligomerization of KIF5A, abolishing autoinhibition and increasing cargo transport (49–52). Consequently, aberrant oligomerization and excess motor activity of KIFs might be common triggers of neurodegeneration. These perspectives indicate the importance of precise motor regulation in maintaining neuronal function. Understanding the molecular mechanisms underlying endogenous KIF1A/UNC-104 dimerization will provide insights for identifying potential therapeutic targets.

### Limitation of this study

We have not determined as to how a single point mutation can cause dimerization of the entire length of DmUNC104, despite the many intramolecular interactions in the full-length DmUNC-104. The mechanism by which the wild type forms dimers is also unknown. *In vitro* reconstitutions using previously identified cargo adaptors such as DENN, Liprin-α and ARL-8 are required for a comprehensive understanding (42, 53, 54).

## Experimental procedures

### Worm experiments

*C. elegans* strains were maintained as described previously (55). N2 wild-type worms and OP50 feeder bacteria were obtained from the *C. elegans* genetic center (CGC) (Minneapolis, MN, USA). *arl-8(wy271)* was described previously (42, 43). Strains used in this study are described in ***supplementary table S2***.

### Genome editing

*unc-104(R551H)* mutation was introduced by Alt-R® CRISPR/cas9 system (Integrated DNA Technologies). Alt-R® CRISPR-Cas9 tracrRNA (#1072533), crRNA for *unc-104* and Alt-R® *S.p.* Cas9 (#1081058) were purchased from Integrated DNA Technologies. The target sequence was 5’-ACAGGATCTAGAGTTATTCT -3’ (Supplementary Figure S1)

The repair template was

5’-ATAAATGGAAAACAAGTGACAACTCCTACTGTATTACACACAGGATCCCACGTTATCCTAGGTGAACATCACGTTTTCCGATATAATGATCCACAGGAAG-3’

Injection mix was prepared as described with a slight modification (39). We excluded *rol-6* marker because successful injections should produce mutants with *unc* phenotypes due to deletion mutations of *unc-104* caused by repair failures. 3-days later after the injection, we selected a few plates that contain strong *unc* worms. Then, *unc* mutant worms as well as superficially *wild-type* worms were singled from these plates. 7 days later, plates that contain *unc-104(R551H)* allele was selected by genomic PCR and BamHI (NEB) digestion. 30% of plates contained either *unc-104(R551H)* heterozygotes or homozygotes.

### Plasmids

cDNA of Drosophila *unc-104* (*Dmunc-104*) was described previously (33). *Dmunc-104* cDNA fragments were amplified by PCR and cloned into the pAcebac1 vector with C-terminal sfGFP and 2xStrep-tag II (25) by Gibson assembly (56). G618R, R562H(*bris*), V6M, A255V and R348G mutations were introduced in the pACEBac1_Dmunc-104::sfGFP::Strep-tagⅡwith the following primers. The sequence of the plasmids were verified by Sanger sequencing. Plasmids used in the study are summarized in ***supplementary table S1*.**

*G618R*_F, gcgaattgctcgagaagcaacgcattgatctaaaagc

*G618R*_R, gcttttagatcaatgcgttgcttctcgagcaattcgc

*R562H*_F, cttaagaccggttctcacgtgatcctcggaaag

*R562H*_R, ctttccgaggatcacgtgagaaccggtcttaag

*V6M*_F, gcggaattcatgtcgtcggttaagatggcggtgcgagtgcg

*V6M*_R, cgcactcgcaccgccatcttaaccgacgacatgaattccgc

*A255V*_F, cttggccgggtcggaacgagtagattccactggtgccaaggg

*A255V*_R, cccttggcaccagtggaatctactcgttccgacccggccaag

*R348G*_F, gcacactgcgctatgcggatggtgccaagcaaattgtttgcaagg

*R348G*_R, ccttgcaaacaatttgcttggcaccatccgcatagcgcagtgtgc

### Expression of DmUNC-104 in Sf9 cells

Sf9 cells (Thermo Fisher Scientific) were maintained in Sf900^TM^ II SFM (Thermo Fisher Scientific) at 27°C. DH10Bac (Thermo Fisher Scientific) were transformed with the plasmids to generate bacmid. To prepare baculovirus, 1 × 10^6^ of Sf9 cells were transferred to each well of a tissue-culture treated 6 well plate. After the cells attached to the bottom of the dishes, ∼5 μg of bacmid were transfected using 5 μL of TransIT^®^-Insect transfection reagent (Takara Bio Inc.). 5 days after initial transfection, the culture media were spun at 15,000 × g for 1 min to obtain the supernatant (P1). For protein expression, 400 mL of Sf9 cells (2 × 10^6^ cells/mL) were infected with 200 µL of P1 virus and cultured for 65 h at 27°C. Cells were harvested and stocked at -80°C.

### Purification of recombinant proteins

Sf9 cells were resuspended in 25 mL of lysis buffer (50 mM HEPES-KOH, 150 mM KCH3COO, 2 mM MgSO4, 10% glycerol, pH 7.5) along with 0.1 mM ATP, 1 mM PMSF, 0.1 mM AEBSF, 0.1 µM Aprotinin, 5 µM Bestatin, 2 µM E-64, 2 µM Leupeptin and 1 µM Pepstatin A. Then 0.5% Triton X-100 were added to lyse the cells. After incubating on ice for 10 min, lysates were cleared by centrifugation (50,000 × g, 20 min, 4°C) and subjected to affinity chromatography described below.

Lysate was loaded on 5 ml of Streptactin-XT resin (IBA Lifesciences, Göttingen, Germany) and passed through the resin by gravity flow. The resin was washed with 30 ml of lysis buffer. Protein was eluted with 25 ml of elution buffer (50 mM HEPES-KOH, 150 mM KCH3COO, 2 mM MgSO4, 10% glycerol, 100 mM D-biotin, pH 7.5). Eluted protein was concentrated using an centrifugal filters with MWCO 50K (AS ONE Corporation #4-2669-05), flash frozen in liquid nitrogen and stored at - 80°C. SEC was performed using an NGC chromatography system (Bio-Rad) equipped with BioSep-SEC-s4000, particle size 5 mm, pore size 500A°, 7.8 mm ID x 600 mm column (Phenomenex) equilibrated in GF150 buffer (25 mM HEPES-KOH, 150 mM KCl, 2 mM MgCl2, pH 7.2). Fractions were analyzed by SDS-PAGE and visualized using the stain-free protocol (Bio-Rad) with ChemiDoc^TM^ Imaging System (Bio-Rad). Peak fractions were collected and concentrated using an Amicon Ultra-0.5 (Merck). Protein concentration was determined by the absorbance of sfGFP at 488 nm using NanoDrop^TM^ (Thermo Fisher Scientific). Proteins at 1-2 µM (as dimers) were flash frozen in liquid nitrogen and stored at -80℃ for further analysis.

### Mass photometry

Proteins were thawed and diluted to the final concentration of 10 nM in GF150 buffer (concentration assumed as dimers). Mass photometry was performed using a Refeyn OneMP mass photometer (Refeyn Japan, Kobe, Japan) and Refeyn AcquireMP version 2.3 software, with default parameters set by Refeyn AcquireMP. Bovine serum albumin (BSA) was used as a control to determine the molecular weight. The results were subsequently analyzed using Refeyn DiscoverMP version 2.3, and graphs were prepared to show the distribution of molecular weight.

### TIRF single-molecule motility assays

For preparing microtubules for TIRF assays, fresh pig brains were obtained from the Shibaura Slaughterhouse in Tokyo. Tubulin was purified from the brain as described (57). Tubulin was labeled with Biotin-PEG_2_-NHS ester (Tokyo Chemical Industry, Tokyo, Japan) and AZDye647 NHS ester (Fluoroprobes, Scottsdale, AZ, USA) as described (58). To polymerize Taxol-stabilized microtubules labeled with biotin and AZDye647, 30 μM unlabeled tubulin, 1.5 μM biotin-labeled tubulin and 1.5 μM AZDye647-labeled tubulin were mixed in BRB80 buffer supplemented with 1 mM GTP and incubated for 15 min at 37°C. Then, an equal amount of BRB80 supplemented with 40 μM taxol was added and further incubated for 30 min. The solution was loaded on BRB80 supplemented with 30% sucrose and 20 μM taxol and ultracentrifuged at 100,000 g for 5 min at 30°C. The pellet containing polymerized microtubules was resuspended in BRB80 supplemented with 20 μM taxol and used for TIRF assays. Glass chambers were prepared by acid washing as previously described (59). Glass chambers were coated with PLL-PEG-biotin (SuSoS, Dübendorf, Switzerland). Polymerized microtubules were flowed into streptavidin adsorbed flow chambers and allowed to adhere for 5–10 min. Unbound microtubules were washed away using assay buffer (90 mM HEPES-KOH, 50 mM KCH_3_COO, 2 mM Mg(CH_3_COO)_2_, 1 mM EGTA, 10% glycerol, pH 7.4) supplemented with 0.1 mg/ml biotin–BSA, 0.2 mg/ml kappa-casein, 0.5% Pluronic F127, 2 mM ATP, and an oxygen scavenging system composed of PCA/PCD/Trolox. Purified motor protein was diluted to indicated concentrations in the assay buffer and flowed into the glass chamber. An ECLIPSE Ti2-E microscope equipped with a CFI Apochromat TIRF 100XC Oil objective lens (1.49 NA), an Andor iXion life 897 camera and a Ti2-LAPP illumination system (Nikon, Tokyo, Japan) was used to observe single molecule motility. NIS-Elements AR software ver. 5.2 (Nikon) was used to control the system.

### Statistical analyses and graph preparation

Statistical analyses were performed using Graph Pad Prism version 10. Statistical methods are described in the figure legends. Graphs were also prepared using Graph Pad Prism version 10. Alignment of amino acid sequences were performed using Clustal WS with default settings of Jalview (Clustal W and Clustal X version 2.0) (60).

## Data availability

Plasmids used for protein expression have been deposited at Addgene.

## Supporting information

This article contains supporting information, including Figures S1 and S2, Tables S1 and S2.

## Acknowledgments

We thank Dr. Thomas Schwarz (Harvard Medical School) for generously providing the cDNA of Drosophila *unc-104.* We thank Atsushi Nakagawa and Jiye Wang (Osaka University) for help with mass photometry. During the preparation of this manuscript, the authors used ChatGPT in order to check English grammar and improve English writing. After using this tool, the authors reviewed and edited the content as needed and take full responsibility for the content of the publication.

## Author Contributions

S.N. and K.C. designed research; S.N., T.W. and K.C. performed research; S.N. and K.C. analyzed data; S.N. and K.C. wrote the paper.

## Funding and additional information

SN was supported by JSPS KAKENHI (23H02472 and 22H05523), the Naito foundation and the Uehara Memorial foundation. KC was supported by JSPS KAKENHI (22K15053), the Naito foundation and MEXT (Ministry of Education, Culture, Sports, Science and Technology) Leading Initiative for Excellent Young Researchers (JPMXS0320200156). Some worm strains and OP50 were obtained from the CGC. This work was performed under the Collaborative Research Program of Institute for Prorein Research, Osaka University, CR-23-02.

## Abbreviations and nomenclature

KIF: kinesin superfamily protein
MT: microtubule
UNC: uncoordinated
*C. elegans*: Caenorhabditis elegans
DCV: dense core vesicle
KAND: KIF1A-associated neurological disorder
MD: motor domain
CC: coiled-coil domain;
NC: neck coiled-coil
FHA: forkhead- associated domain
MBS: membrane-associated guanylate kinase homolog (MAGUK)-binding stalk domain
MATH: meprin and TRAF homology domain
PH: pleckstrin homology domain
IMAC: immaculate connections
KLP: kinesin-like protein
ARL: arf-like small guanosine triphosphatase
TIRF: total internal reflection fluorescent
DLS: dynamic light scattering;
SEC: size exclusion chromatography
EMS: ethyl methanesulfonate
Brp: Bruchpilot
AEBSF: 4-benzenesulfonyl fluoride hydrochloride
PMSF: phenylmethylsulfonyl fluoride
sfGFP: superfolder GFP
AMPPNP: Adenylyl imidodiphosphate
Kd: dissociation constant
M.W.: molecular weight

---

**Figure S1.**
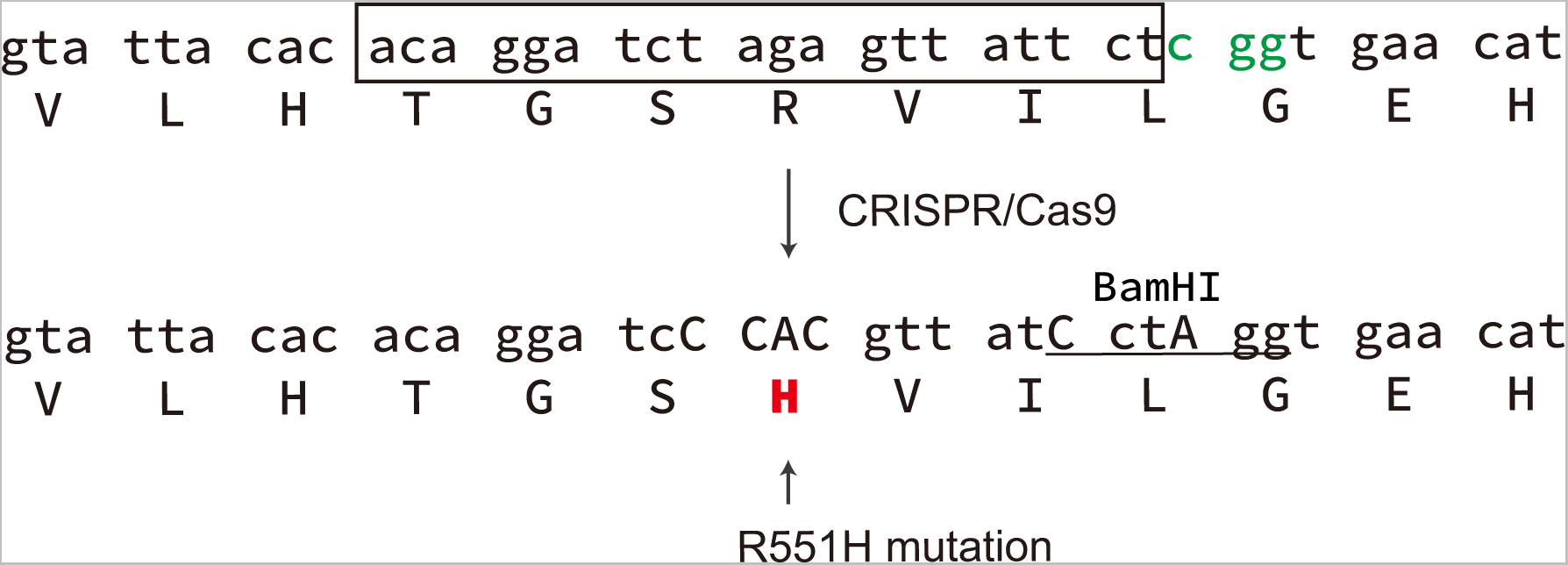
Schematic drawing showing the genome editing to introduce R551H mutation. The genome sequence of *C. elegans* unc-104 encoding CeUNC-104(R551) (upper) and edited sequence (lower) are shown. Green font indicates PAM sequence. The boxed sequence is the target sequence for Cas9. Uppercase letters indicate edited bases. Note that a BamHI site was introduced to check the editing.

**Figure S2.**
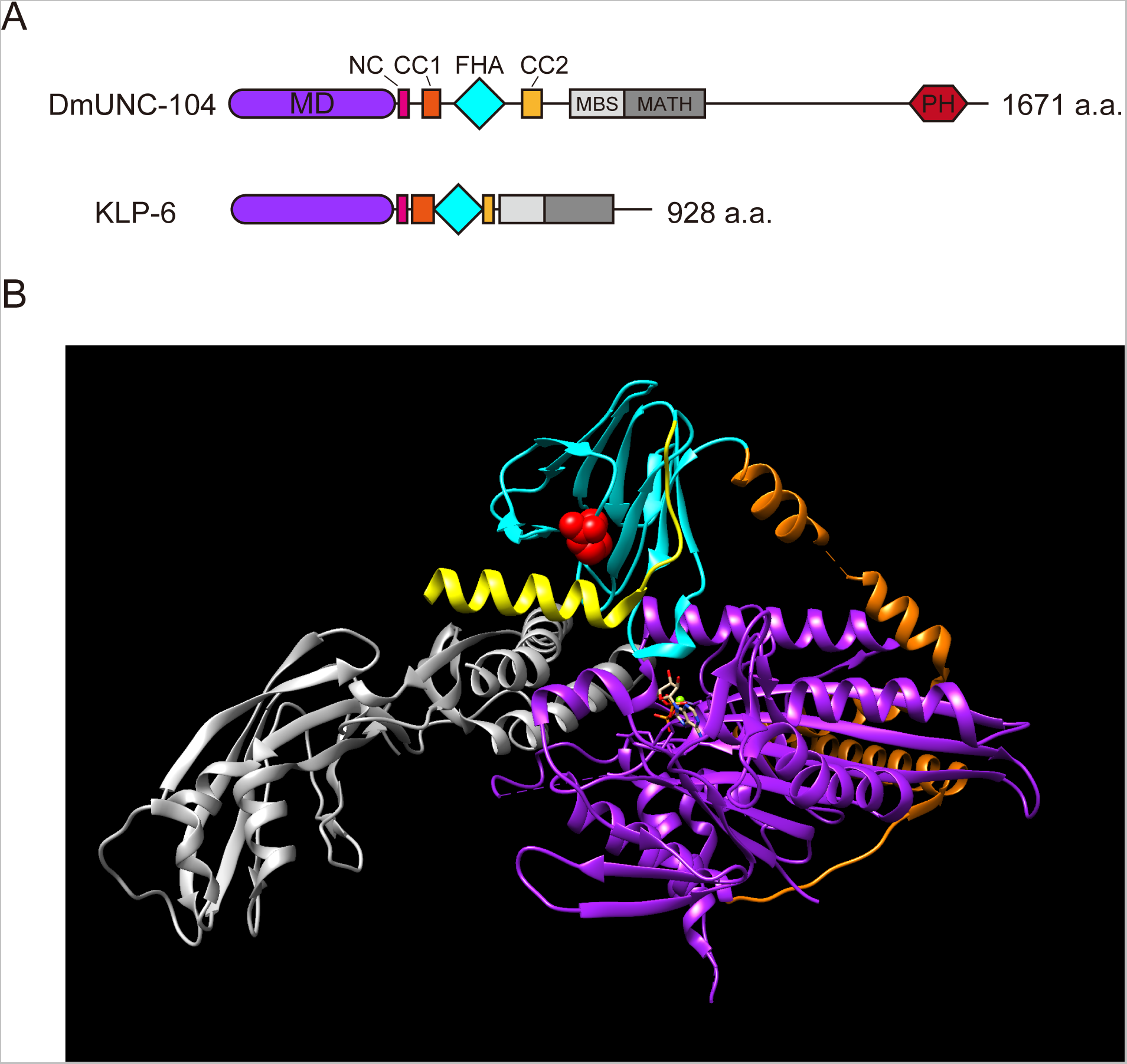
The position of R562 corresponding residue on the full-length KLP-6 structure. **(A)** Schematic diagram of the domain architecture of *Drosophila* UNC-104 (DmUNC-104) and *C. elegans* KLP-6 (KLP-6). MD: motor domain, NC: Neck coiled-coil domain, CC1: Coiled-coil 1 domain, FHA: forkhead-associated domain, CC2: Coiled-coil 2 domain, MBS: membrane-associated guanylate kinase homolog (MAGUK)-binding stalk domain, MATH: meprin and TRAF homology domain, PH: Pleckstrin homology domain. **(B)** The full-length KLP-6 structure (PDB accession no. 7WRG; Wang et al., 2022) is shown with distinct domains highlighted. MD (purple), NC (orange), CC1 (orange), FHA domain (cyan), CC 2 (yellow), MBS (gray) and MATH (gray) are shown. The R538 residue, corresponding to R562 of DmUNC-104, is shown by red.

**Supplementary Table S1.** Plasmid List.

| Plasmid Name | Description | Comment | Source |
| --- | --- | --- | --- |
| pSN539 | pACEBac1 Dmunc-104-sfGFP-2xStrepII | Figure 1 | This study |
| pKC31 | pACEBac1 Dmunc-104(R562H(bris))-sfGFP-2xStrepII | Figure 2 | This study |
| pKC28 | pACEBac1 Dmunc-104(G618R)-sfGFP-2xStrepII | Figure 4 | This study |
| pKC50 | pACEBac1 Dmunc-104(V6M)-sfGFP-2xStrepII | Figure 5 | This study |
| pKC51 | pACEBac1 Dmunc-104(A255V)-sfGFP-2xStrepII | Figure 5 | This study |
| pKC52 | pACEBac1 Dmunc-104(R348G)-sfGFP-2xStrepII | Figure 5 | This study |

**Supplementary Table S2.** Strain List.

| Strain Name | Genotype | Comment | Source |
| --- | --- | --- | --- |
| TV1229 | *wyIs85* V | Figure 3C | Klassen et al., 2007 |
| OTL81 | *arl-8*(wy271)IV; *wyIs85* V | Figure 3D | Wu et al., 2013 |
| OTL220 | *unc-104(R551H)* II; *arl-8*(wy271)IV; *wyIs85* V | Figure 3E | This study |

